# *ENHANCED GRAVITROPISM 2* encodes a STERILE ALPHA MOTIVE containing protein that controls root growth angle in barley and wheat

**DOI:** 10.1101/2021.01.23.427880

**Authors:** Gwendolyn K. Kirschner, Serena Rosignoli, Isaia Vardanega, Li Guo, Jafargholi Imani, Janine Altmüller, Sara G. Milner, Raffaella Balzano, Kerstin A. Nagel, Daniel Pflugfelder, Cristian Forestan, Riccardo Bovina, Robert Koller, Tyll G. Stöcker, Martin Mascher, James Simmonds, Cristobal Uauy, Heiko Schoof, Roberto Tuberosa, Silvio Salvi, Frank Hochholdinger

## Abstract

The root growth angle defines how roots grow towards the gravity vector and is among the most important determinants of root system architecture. It controls water uptake capacity, nutrient use efficiency, stress resilience and as a consequence yield of crop plants. We demonstrated that the *egt2* (*enhanced gravitropism 2*) mutant of barley exhibits steeper root growth of seminal and lateral roots and an auxin independent higher responsiveness to gravity compared to wild type plants. We cloned the *EGT2* gene by a combination of bulked segregant analysis and whole genome sequencing. Subsequent validation experiments by an independent CRISPR/Cas9 mutant allele demonstrated that *egt2* encodes a STERILE ALPHA MOTIF domain containing protein. *In situ* hybridization experiments illustrated that *EGT2* is expressed from the root cap to the elongation zone. Subcellular localization experiments revealed that *EGT2* localizes to the nucleus and cytoplasm. We demonstrated the evolutionary conserved role of *EGT2* in root growth angle control between barley and wheat by knocking out the *EGT2* orthologs in the A and B genomes of tetraploid durum wheat. By combining laser capture microdissection with RNA-seq, we observed that seven expansin genes were transcriptionally downregulated in the elongation zone. This is consistent with a role of *EGT2* in this region of the root where the effect of gravity sensing is executed by differential cell elongation. Our findings suggest that *EGT2* is an evolutionary conserved regulator of root growth angle in barley and wheat that could be a valuable target for root-based crop improvement strategies in cereals.

**Significance Statement:** To date the potential of utilizing root traits in plant breeding remains largely untapped. In this study we cloned and characterized the *ENHANCED GRAVITROPISM2* (*EGT2*) gene of barley that encodes a STERILE ALPHA MOTIF domain containing protein. We demonstrated that *EGT2* is a key gene of root growth angle regulation in response to gravity which is conserved in barley and wheat and could be a promising target for crop improvement in cereals.

## Introduction

The increase of human population and climate change are major challenges to food security (1, 2). A number of studies proposed to modify root system architecture to improve water and nutrient use efficiency, crop yield and resilience to stress episodes (3, 4). Among the most important determinants of root system architecture is the root growth angle, i.e. the angle in which roots grow towards the ground.

Increased response to gravity, or hypergravitropism, and thereby a steeper root growth angle was shown to be associated to improved drought resistance in rice, probably by increased access to deep-soil water (5). At the same time, a deeper root system facilitates the uptake of N and other mobile nutrients which are more abundant in deeper soil layers (6). Root gravitropism is regulated by sensing the gravitropic stimulus and subsequent differential cell elongation to enable root growth towards the gravitropic vector. Removing the root cap mechanically or genetically substantially diminishes the gravitropic response (7–9) suggesting that gravity sensing occurs primarily in the root cap. However, there is evidence for a sensing site outside the root cap, located in the elongation zone (10, 11). There are different hypotheses on how the cells sense gravity, with the prevailing idea that the starch-containing plastids in the root cap act as statoliths and settle in response to gravity. In doing so, they trigger a signaling cascade, either by mechanosensitive channels or by direct protein interaction, on the organelle surface (12–14). This signaling pathway ultimately leads to a rearrangement of auxin export carriers and thereby to a reorganization of the auxin maximum in the root tip (15). At the same time, changes of pH in the root cap and an asymmetrical change of pH in the upper and lower side of the root meristem and elongation zone occur (16, 17). This finally leads to an increased elongation of the cells on the side averted to the gravity vector in the elongation zone of the roots, so that the roots grow downwards (18). To date, only single components of the signaling cascade regulating root gravitropism have been unraveled. Examples include the actin-binding protein RICE MORPHOLOGY DETERMINANT that localizes to the surface of statoliths in rice root cap cells and controls the root growth angle in response to external phosphate (19). Another protein involved in gravitropism is the membrane-localized ALTERED RESPONSE TO GRAVITY1 in Arabidopsis, which is expressed in the root cap and is involved in the gravity-induced lateral auxin gradient (20). Both proteins seem to function in signaling immediately after gravity sensing in the root cap. In contrast, rice *DEEPER ROOTING1* (*DRO1*), acts as early auxin response gene later in the gravitropic signaling. The *DRO1* gene encodes for a plasma membrane protein that is expressed in the root meristem and was identified because of its influence on the root growth angle (5). The role of DRO1 may not be conserved in primary roots of different plant species, since the Arabidopsis homolog does not affect the gravitropic response of the primary root but influences the growth angle of the lateral roots (21).

Barley (*Hordeum vulgare*) is the world’s fourth most important cereal crop in terms of grain production, after wheat (*Triticum aestivum*), rice (*Oryza sativa*) and maize (*Zea mays*) (2017, http://faostat.fao.org). It is cultivated over a broad geographical area because it can adapt to a wide range of climatic conditions and is therefore an excellent model to study responses to climate change (22). In this study we used a forward-genetics approach to clone *ENHANCED GRAVITROPISM2* (*EGT2*), a novel gene involved in barley root gravitropic response and whose effect is conserved in wheat. *EGT2* encodes a STERILE ALPHA MOTIF domain containing protein and likely acts in a regulatory pathway that counteracts the auxin-mediated positive gravitropic signaling pathway.

## Results

### The *egt2-1* mutant shows a steeper root growth correlated with an enhanced gravitropic response

The *egt2-1* mutant was discovered in a sodium-azide mutagenized population of the barley cultivar Morex based on the hypergravitropic growth of its seminal root system in paper rolls and shown to be inherited as a monogenic recessive Mendelian locus (23–25). We investigated the phenotype in more detail in 2-D rhizoboxes, in which the plants grow vertically on flat filter paper. While in Morex wild type the seminal roots grow in a shallow angle towards the gravity vector and cover a larger area, the seminal roots in the *egt2-1* mutant grow steeply down (Figure 1A, Supplementary Figure 1A). This phenotype was consistent in plants grown in soil-filled rhizotrons and pots, the latter visualized by magnetic resonance imaging (MRI) (Figure 1B, C, Supplementary Figure 1E, F, G). Furthermore, the lateral roots arising from the seminal roots also displayed a highly increased growth angle (Figure 1B, E, Supplementary Figure 1A, H). Apart from the increased root growth angle, we did not detect any other aberrant root phenotypes, neither a changed number of seminal roots nor a difference in root length (Supplementary Figure 1B, C). To further investigate the reason for the steep root phenotype, we tested the responsiveness of the root system to gravity. After rotation by 90°, we monitored the angle of the root tips over time (Figure 1F, G). Roots of the *egt2-1* mutant bent much faster and stronger than wild type roots, approaching 90° after 3 days compared to just 30° in wild type roots (Figure 1G). Root growth rate, however, was not altered (Supplementary Figure 1D). We concluded therefore that the steep root angle of the *egt2-1* mutant was likely caused by a higher responsiveness to gravity. Since gravity sensing and signal transduction was shown to take place in the root cap and meristem (16–18), we compared the root cap and meristem by microscopy and measuring the root meristem size, but we did not discover significant differences (Supplementary Figure 2).

**Figure 1:**
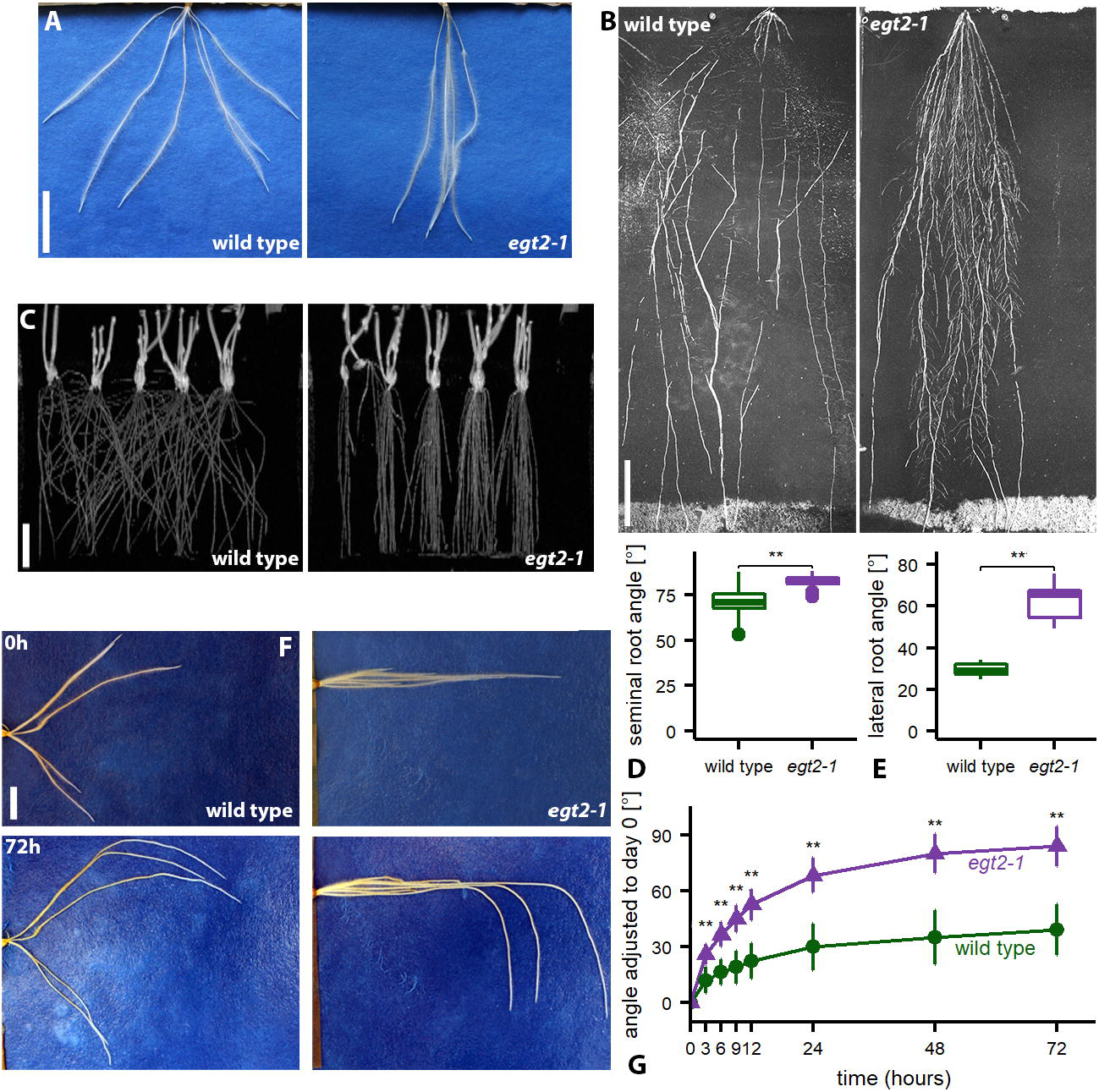
Root phenotype of *egt2-1*. **A** Wild type and *egt2-1* roots grown on germination papers, 7 days after germination (DAG). Scale bar: 2 cm. **B** Wild type and *egt2-1* roots grown in rhizotrons 26 DAG; scale bar: 10 cm. **C** MRI pictures of wild type and *egt2-1* plants grown in soil 3 DAG. Scale bar: 4 cm. **D** Root angle of seminal roots 7 DAG; n = 40 per genotype in one experiment; two-tailed t-test, ** *p* <0.01. **E** Lateral root angle 14 DAG; n = 8-9 per genotype in two independent experiments; two-tailed t-test, ** *p* <0.01. **F** Wild type and *egt2-1* roots after rotation (time point 0) at indicated time points. Scale bar:1 cm. **G** Root tip angle after rotation; plants 5 DAG were rotated by 90° (time 0) and the root tip angle was measured over time; n = 38 per genotype in three independent experiments; the two genotypes were compared between each other at the respective time points by a two-tailed t-test, ** *p* <0.01; standard deviation is depicted; to account for the different starting angles of the roots, all measurements were normalized to the starting angle of the roots at time 0.

### Auxin response is unaffected in *egt2-1* mutants

It was shown before that the phytohormone auxin is involved in gravitropic response signaling (15) and that auxin transport inhibitors or external supply of auxin influences the reaction of roots to rotation (26). To analyze if the *egt2-1* mutant is sensitive to manipulation of the auxin state in the roots, we treated wild type and mutant with auxin or auxin transport inhibitors and recorded the reaction to 90° rotation. Application of the endogenous auxin Indole-3-acetic acid (IAA) as well as the synthetic auxin analog 1-Naphthaleneacetic acid (NAA) led to a faster, albeit non-significant, reaction in both wild type and *egt2-1* mutant immediately after the rotation (Supplementary Figure 3A, C). Treatment with 2,4-Dichlorophenoxyacetic acid (2,4D), an auxin analog which cannot be transported by auxin efflux transporters, caused a significantly faster response in both wild type and *egt2-1* mutant (Supplementary Figure 3E). Treatment with the auxin transport inhibitor 1-N-Naphthylphthalamic acid (NPA) on the other hand, decreased the reaction to rotation significantly in wild type and mutant to a similar degree (Supplementary Figure 3G). Recently, cytokinin was described as anti-gravitropic signal in lateral roots, therefore we included cytokinin in our study (27). However, we found no impact of cytokinin on the root angle after rotation (Supplementary Figure 3I). In summary, we demonstrated that *egt2-1* reacts to auxin treatments to the same degree as the wild type and we conclude that the mutation in *egt2-1* does not disrupt the major auxin signaling pathways. This notion is consistent with the results of a tissue-specific RNA-seq analysis of wild-type and *egt2-1* seminal roots where we did not find any auxin related genes among the differentially expressed genes (see results below).

### *EGT2* encodes a Sterile alpha motif domain-containing protein

In order to map and clone the *EGT2* gene, a SNP-based bulked-segregant analysis (BSA) was carried out using an F_2_-population derived from the cross between the hypergravitropic *egt2*-*1* carrying line TM2835 (in Morex background) and cv. Barke, the latter showing a typical wild-type, shallow root architecture. *egt2* was mapped to a 312 Mbp interval on the short arm of chromosome 5H (Figure 2A), between markers *SCRI_RS_222345* and *SCRI_RS_13395* (Table S1). Subsequently, TM2835 was subjected to whole genome sequencing, which led to the identification of seven genes within the *egt2* interval and which carried missense, splice site or stop-codon gain mutations when compared with wild type Morex sequence (Table S2). Among these was a gene encoding for a 252 amino acid sterile alpha motif (SAM) domain-containing protein (*HORVU5Hr1G027890* (24) or *HORVU.MOREX.r2.5HG0370880.1* (28)) with a mutation (G447A) leading to a premature stop codon at the beginning of the functional domain (W149*) (Figure 2B, Supplementary Figure 4A and B) (28). Apart from the SAM domain, no other functional domains were predicted (29). The sequence of the SAM domain between EGT2, and previously described SAM domains of other plant species is highly conserved (Supplementary Figure 4B, Supplementary Figure 5) (30, 31).

**Figure 2:**
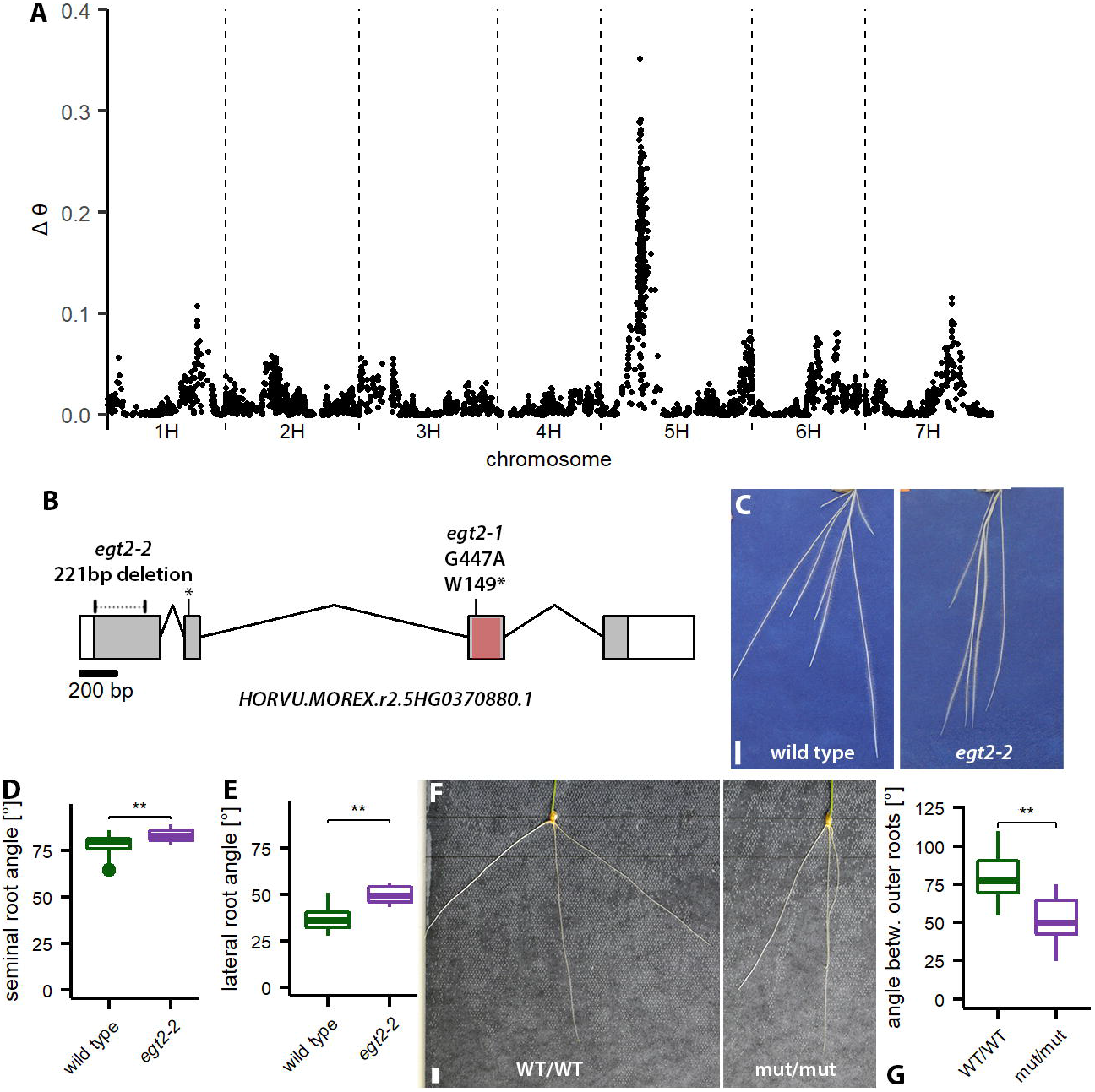
*EGT2* encodes a STERILE ALPHA MOTIF protein. **A** Association of SNP markers with seminal root angle across the barley genome as established by bulk segregant analysis (BSA) in the F_2_ cross TM2835 (*egt2-1*, hypergravitropic roots) × cv. Barke (wt roots). Y axis reports Δθ, an index accounting for the difference in allele-specific fluorescence signal between the two BSA DNA bulks, per SNP. **B** Gene structure of *EGT2* (HORVU.MOREX.r2.5HG0370880.1) with mutations in *egt2* (*egt2-1*: G to A transition and *egt2-2*_deletion); exons depicted as gray box, introns depicted by lines, UTRs depicted as white boxes; red box indicates the sequence encoding for the SAM domain. **C** Exemplary pictures of wild type (cv. Golden Promise) and mutant *egt2-2* roots 7 DAG. Scale bar: 2 cm. **D** Seminal root angle of wild type (cv. Golden Promise) and mutant *egt2-2* 7 DAG; n = 15-17 in two independent experiments. **E** Root angle of lateral roots 14 DAG; n = 16-18 in two independent experiments; two-tailed t-test, * *p* <0.05, ** *p* <0.01. **F** Exemplary pictures of wheat wild type (WT/WT) and *egt2* (mut/mut) roots, 7 days after germination (DAG). Scale bar: 1 cm. **G** Root angle between 2^nd^ and 3^rd^ seminal root of wild type (WT/WT) and *egt2* (mut/mut) wheat seedling at 7 DAG; n = 18 and 39 for wt and mutant, respectively. Wheat plants were derived from two independent segregating populations.

To validate *HORVU.MOREX.r2.5HG0370880.1* as *EGT2*, we used CRISPR/Cas9 to create an additional mutant allele (*egt2-2*) in the barley cv Golden Promise. We targeted two sites in the 5’UTR and exon 1, separated by 196 bp, and recovered a 221 bp deletion leading to a premature stop codon and a truncated 75 amino acid protein (Figure 2B, Supplementary Figure 4A). We analyzed the root phenotype of the homozygous T_1_-line and determined a significantly higher root angle of both seminal and lateral roots in the mutant in comparison to the wild type (Figure 2C, D, E, Supplementary Figure 4D). Hence, we confirmed that the altered root angle phenotype of *egt2-2* is caused by a truncation of *HORVU.MOREX.r2.5HG0370880.1*. Like in the *egt2-1* mutant in Morex background, the root length of *egt2-2* was similar to the wild type (Supplementary Figure 4C). The reaction of the *egt2-2* roots after rotation was faster than in the wild type, however, not statistically significant (Supplementary Figure 4E). It is notable that Golden Promise and Morex differ in seminal root angle growth although they both carry a wild-type *EGT2* allele (compare Figure 1A and E and Figure 2C and D). Additionally, the re-orientation of the roots after rotation occurs much faster in wild type Golden Promise than in Morex (compare Figure 1H and Supplementary Figure 4E). Thus, other genetic factors influence the root growth angle in addition to *EGT2*.

To further validate the function of the *EGT2* gene, we identified mutant lines carrying premature stop codons from a sequenced mutant population of tetraploid wheat (32). We combined mutations in the two durum wheat *EGT2* orthologs (homologs on A and B genomes) to generate complete *egt2* knockout lines. These double mutants showed narrower seminal root growth angle in rhizoboxes compared with the sibling lines carrying wildtype alleles in both homologs (Figure 2F, G). Similar to barley, the number and length of seminal roots was unaffected in 7-day old seedlings (Supplemental Figure 4G, H, I).

### *EGT2* is expressed in the whole root tip and localizes to cytoplasm and nucleus

To survey the spatial expression patterns of *EGT2* in roots we performed RNA *in situ* hybridization experiments. *EGT2* is expressed in the whole root tip, including the root cap, meristem and elongation zone (Figure 3A). We confirmed this expression pattern by RNAseq data, where we found *EGT2* expressed in root cap, meristem and elongation zone and a transcriptional downregulation only in the elongation zone of the mutant (Figure 3B). qRT-PCR analysis of combined root cap, meristem and elongation zone and the differentiation zone confirmed the expression in all these tissues in wild type and mutant (Figure 3C). Transient expression in *Nicotiana benthamiana* leaves revealed that EGT2 localizes to cytoplasm and nucleus (Figure 3D).

**Figure 3:**
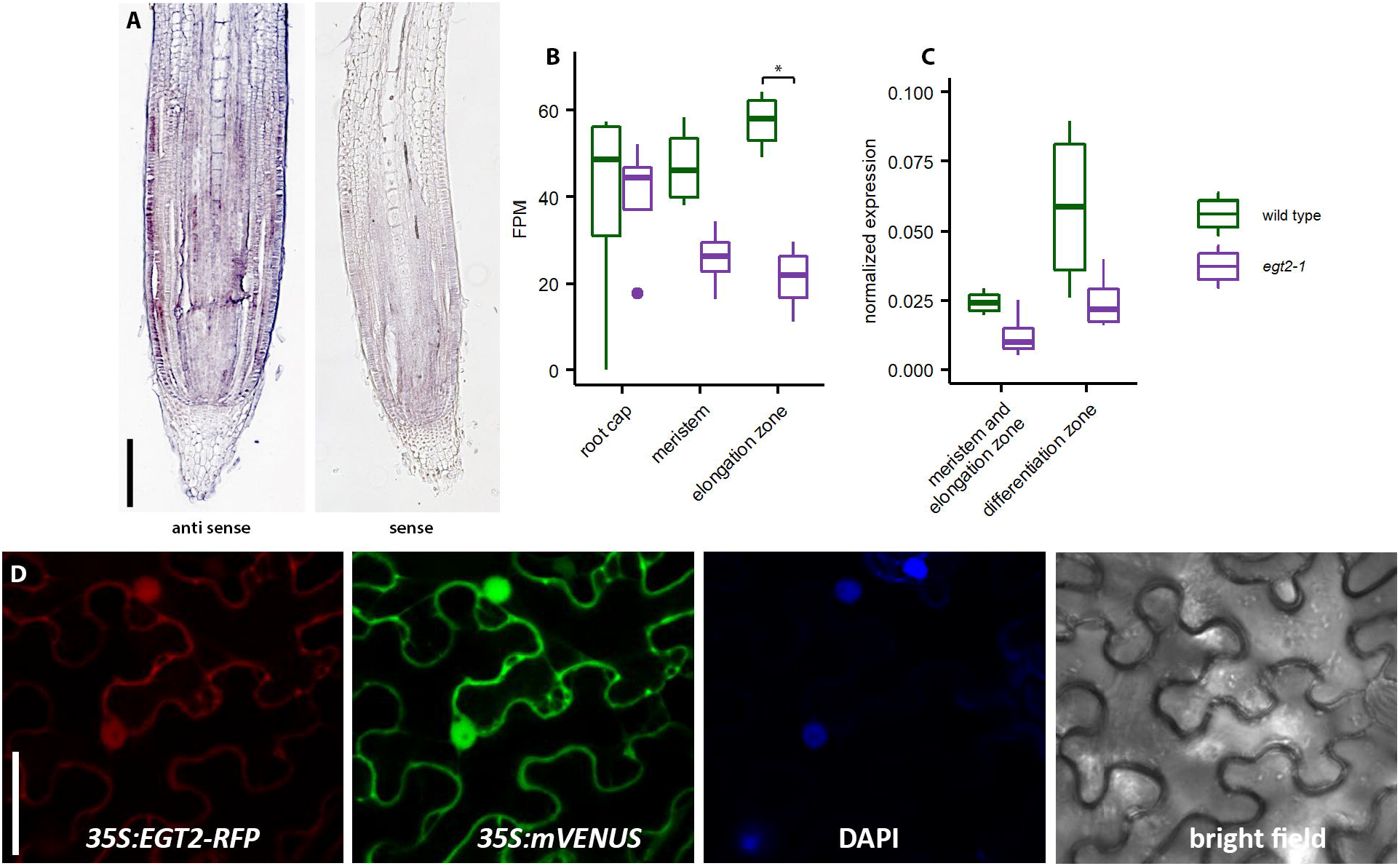
Expression of *EGT2*. **A** RNA *in situ* hybridization of *EGT2*; negative controls (sense probes) are shown on the right. Scale bar: 200 μm. **B** FPM normalized values for *EGT2* expression from the RNAseq dataset in the respective tissues (compare tissue Figure 4D); \**p*_*adj*_ <0.05. **C** qRT-PCR of *EGT2* expression; normalized to *tubulin*; two-tailed t-test does not display significant differences. **D** EGT2 localization in tobacco. Scale bar: 50 μm.

### In the *egt2-1* mutant, cell wall related processes are affected in the elongation zone

To analyze the effect of the mutation in *EGT2* on the root transcriptome, we isolated RNA from different root tissues. For this, we applied laser capture microdissection to specifically separate root cap, meristem and parts of the elongation zone from wild type and *egt2-1* seminal roots. This allowed us to differentiate between gravity sensing (root cap), signal transduction (meristem) and signal execution (elongation zone) (Figure 4B). To this end, we selected the most vertically grown seminal roots in both genotypes that displayed a similar root growth angle (Figure 4A). By doing so we excluded secondary effects caused by different root growth angles. Moreover, we used roots of similar length to exclude differences in age since the barley seminal roots do not grow out simultaneously (33). We determined the transcriptomic relationships among the two genotypes and three tissues by a principle component analysis (PCA) (Figure 4B). In the PCA, the two principle components PC1 and PC2 explained 82% of the total variance (Figure 4B). The biological replicates per tissue including four wild-type and four mutant samples clustered closely together. This indicates small transcriptomic differences between the genotypes but large differences between the tissues. To identify differentially regulated genes, we computed pairwise contrasts between the genotypes of the respective tissues (FDR <5% and log_2_FC >|1|; see methods) for genes that uniquely mapped to chromosomes 1 to 7 (34). This resulted in 67 differentially regulated genes (DEGs) among all tissues, some of which were shared between all or two tissues (Supplementary Figure 6, Table S4). Strikingly, we found seven genes encoding for expansins downregulated in the elongation zone (Supplementary Figure 6, Supplementary Figure 7). Gene ontology (GO) terms were only assigned to genes downregulated in the elongation zone, all of them related to the term cell wall (Figure 4E). At the same time, this validates our data set, since expansins are expressed in the elongation zone and differentiated root tissue (35). Furthermore, we found that several genes categorized as peroxidase superfamily protein members, upregulated in either the meristem or the elongation zone (HORVU2Hr1G026420, HORVU7HR1G020300, HORVU3Hr1G036820) (Supplementary Figure 6). Differential regulation of a peroxidase superfamily protein-encoding gene was already found in a study in Arabidopsis related to agravitropic mutants (36). Moreover, we found that a gene encoding for calmodulin, a primary plant calcium receptor, downregulated in meristem and elongation zone (HORVU1Hr1G068440) (Supplementary Figure 6). Finally, we found a gene annotated as excocyst complex component 7 upregulated in the meristematic zone. Components of the exocyst are involved in directing exocytotic vesicles to fusion sites on the plasma membrane and might be involved in the distribution of the auxin transporter PINFORMED4 in Arabidopsis (37, 38).

**Figure 4.**
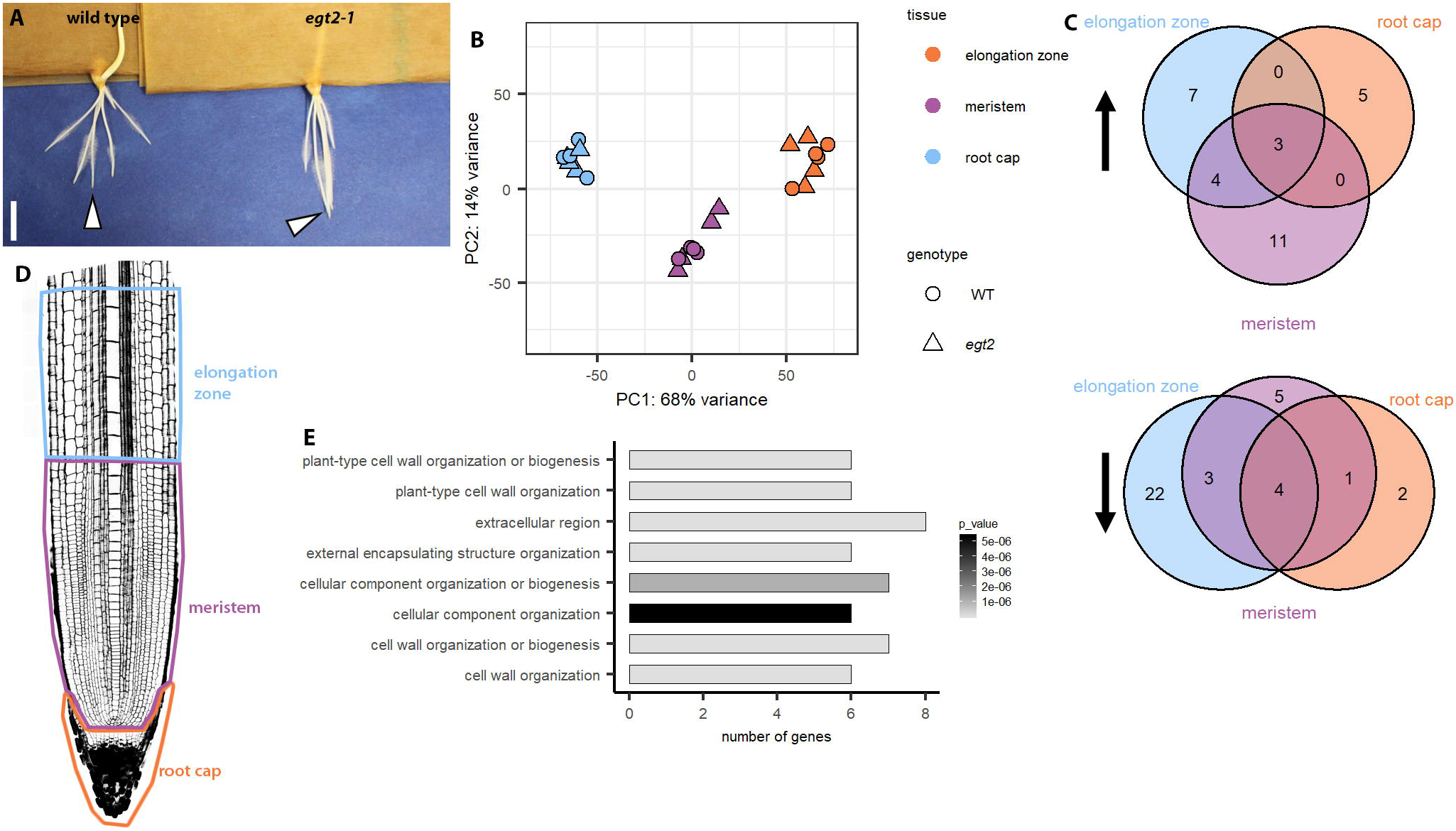
RNAseq reveals differences in cell wall-related processes in the elongation zone. **A** Wild type and *egt2-1* plants 3 DAG used for RNA isolation. Scale bar: 1 cm; arrow heads point to exemplary roots used for RNA isolation (most vertical ones). **B** Principal component analysis (PCA) of the 24 RNA-seq samples of the two genotypes and three tissues; first and second principal components collectively explain 82% of the variance. **C** Venn diagram showing upregulated (upward arrow) and downregulated (downward arrow) differentially expressed genes (DEGs) in the respective tissue. **D** Experimental setup: RNA of root cap, meristem and 900 μm of the elongation zone were isolated. **E** Enriched gene ontology (GO) terms for DEGs downregulated in the elongation zone.

## Discussion

The optimization of root system architecture has been recognized as one of the most important objectives in current breeding programs aimed at increasing resilience and sustainability of crops and agricultural systems (4, 39). Specifically, variation of root growth angle can affect the way roots explore different soil layers, capture nutrients and water and thus can influence drought tolerance, as shown for *DRO1* in rice (4). However, knowledge about genes, gene interactions and regulatory networks in root development is currently limited in all major crops, including cereals.

Here we cloned a novel key regulator of root gravitropism, *ENHANCED GRAVITROPISM2* (*EGT2*), in barley and wheat. Mutations in *EGT2* lead to enhanced gravitropic response and thereby to a steeper root growth angle of seminal and lateral roots. We did not find any other root or shoot morphological trait affected by this mutation, indicating that *EGT2* does not act in an ubiquitous signaling pathway, but rather is specific for root gravitropism. In Arabidopsis, most gravitropic mutants were discovered because they show agravitropic root phenotypes (20, 40). This is probably because a hypergravitropic phenotype is difficult to detect given that the primary root of Arabidopsis grows highly gravitropic in wild type and that Arabidopsis lacks seminal roots (41). In grasses, however, some mutants with hypergravitropic roots were discovered, for instance *vln2* and *rmd*. The villin protein VLN2 facilitates microfilament bundling, while the actin-binding protein RMD links actin filaments with gravity-sensing organelles (19, 26).

The only predicted domain in EGT2 is the SAM domain. In animals, SAM domain containing proteins function as transcription factors, receptors, kinases or ER proteins (30). In plants, the best known protein containing a SAM domain is the transcription factor LEAFY (LFY) which is involved in flower and meristem identity formation. Modelling of Arabidopsis SAM proteins based on structure predictions and LFY characterization suggests that the majority of these proteins are able to form head-to-tail homo- or hetero-oligomers/polymers (30). The close phylogenetic relationship of EGT2 with AtSAM5 (At3g07760) indicates a similar potential of oligomerization for EGT2.

EGT2 is also closely related to WEEP, a SAM domain containing protein that was discovered because of the prominent shoot phenotype in peach tree mutants (31). Peach trees with deletions in *WEEP* show a weeping shoot growth phenotype, thus the branches grow in a wider angle and after gravistimulation by rotation by 90°, the branches do not orient their growth upwards again (31). Therefore, EGT2 and WEEP are likely involved in a similar pathway that regulates gravitropism, however in opposite directions of the plant growth. Bud grafting experiments in peach implied that *WEEP* encodes an autonomous determinant of shoot orientation for each branch, and that no mobile signals from other parts of the plants (like phytohormones) are necessary (31). Furthermore, no difference of auxin or abscisic acid concentration was detected in growing shoots between peach wild type and *WEEP* mutants, nor were genes associated with auxin biosynthesis or perception differentially expressed (31).

Similarly, we did not find any expression changes of genes related to auxin biosynthesis or perception in our transcriptomic comparison between wild type and *egt2-1*. Treatments with auxins or an auxin transport inhibitor confirmed that the *egt2-1* mutant is as sensitive to disturbance of the auxin balance as the wild type (Supplementary Figure 3), indicating that EGT2 works independently of auxin. Nevertheless, the auxin transport pathway could still be affected, as demonstrated for the rice mutant *villin2* (*vln2*) that exhibits a disturbed recycling of the auxin efflux carrier PINFORMED2 (PIN2) and thereby a hypergravitropic root response (26). However, auxin treatment of *pin2* mutants induced a restoration of the phenotype and the mutants were insensitive to auxin transport inhibitors which differs from *egt2-1*. On the other hand, exocyst complex component 7 was transcriptionally upregulated in the *egt2-1* mutant. In Arabidopsis, disturbing the expression of *EXOCYST70A3* by knock-out or overexpression leads to a higher agravitropic response upon auxin efflux inhibitors probably by regulating the PIN4 localization in columella cells and thereby auxin distribution in root tips (37). It is conceivable that disturbance of expression levels in *egt2-1* mutants might lead to a change in PIN localization and thereby a changed signal transduction, however, this hypothesis remains to be tested. The ubiquitous expression of EGT2 in root cap, meristem and elongation zone suggests a participation rather in the signal transduction of gravitropism than in the sensing or differential cell elongation.

Localization of EGT2 in cytoplasm and nucleus is similar to the predicted localization of the homolog AtSAM (30). In a split-ubiquitin yeast two-hybrid screen, AtSAM5 interacted with the calcium dependent protein kinase AtCPK13 (At3G51850) (42), putatively connecting AtSAM5 to calcium signaling pathways. Inhibition of the primary plant calcium receptor calmoldulin was shown to inhibit the response to gravity in Arabidopsis (40). In the *egt2-1* mutant, *Calmodulin 5* is transcriptionally downregulated in meristem and elongation zone, putatively connecting EGT2 with calcium-dependent signal transduction. It is still generally unknown, however, which role calcium plays in gravitropic signaling.

If we hypothesize a role for EGT2 in signal transduction, we would expect downstream targets in the elongation zone, where the effect of gravity sensing is executed by differential cell elongation (18). Here, we found a striking number of expansin genes transcriptionally downregulated (Supplementary Figure 6). Expansins are known as acid-induced cell wall loosening enzymes. However, most studies are based on the activity of bacterial enzymes and the function of expansins in plant cell walls is still unknown (35). Similarly, cell wall-related genes are differentially regulated in *weep* mutant peach trees, and the differentially regulated auxin response genes in *weep* mutants have roles in mediating cell expansion, or modulation of H^+^ transport (31). Besides expansins, we found three genes encoding peroxidase superfamily proteins upregulated in the meristem and elongation zone (Supplementary Figure 6). Downregulation of a peroxidase superfamily protein-encoding gene was already demonstrated in a study in Arabidopsis comparing inflorescence stems of wild type to *scarecrow* and *short root* mutant transcriptomes, which show no gravitropic response to rotation in the shoot (36). Peroxidases catalyze the consumption or release of H_2_O_2_ and reactive oxygen species (ROS). One class of peroxidases functions extracellularly, either for cell wall loosening or cell wall cross-linking (43), and the transcriptional regulation in *egt2* might be related to the regulation of the expansins. Moreover it was shown that ROS work downstream of auxin signaling in root gravitropism, maybe as second messenger (44). Besides the auxin-mediated pathway that directs downward root growth, in Arabidopsis lateral roots, which develop a distinct gravitropic setpoint angle, a pathway was discovered that counteracts the gravitropic bending independently of auxin (27). It is based on cytokinin signaling, indicating that the gravitropic setpoint angle is set by a balance of this pathway and the auxin-mediated positive response to gravity.

Based on the broad expression pattern of *EGT2* throughout root cap, meristem and elongation zone, and the interaction of the Arabidopsis AtSAM5 homolog with CPK13, we can hypothesize that *EGT2* is involved in the signal transduction of gravitropic signaling. The missing interference in auxin-related processes on the transcriptomic level and the susceptibility to auxin treatments implies that EGT2 is not involved in any signal transduction related to changes in auxin levels and/or transport. It is possible that EGT2 acts in a pathway that counteracts the auxin-mediated positive gravitropic signaling pathway, since for the growth in an angle towards the gravity vector, a pathway counteracting the positive reaction to gravity is needed. By knocking it out, the downward growth of the roots would dominate and create the hypergravitropic phenotype.

In summary our results suggest that *EGT2* is an evolutionary conserved check point of seminal and lateral root growth angle in barley and wheat. *EGT2* could be a promising target for root-based crop improvement in cereals.

## Materials and Methods

### Plant material and growth conditions

The *egt2-1* mutation carrying line, TM2835, was derived from sodium azide mutagenesis of the cv. Morex as previously described (24, 25). For growth in rhizoboxes and on agar plates, the seeds were washed in 1.2% sodium hypochlorite for 5 min and rinsed with distilled water. Then they were incubated in darkness at 30 °C over night to induce germination and only germinating seeds were used for further experiments. Growth in rhizoboxes for plant phenotyping and rotation experiments were conducted as described before (46). For phytohormone treatments, plants were grown on half-strength Hoagland solution (47), pH 5.8, supplemented with 0.8% phytagel on square Petri dishes, which were placed in a 45° angle. The plants were grown in growth cabinets (Conviron, Winnipeg, Manitoba, Canada) at 18 °C at night (8 h) and 22 °C at day (16 h). For growth in rhizotrons filled with peat substrate, wild type and *egt2* mutants were grown in the GrowScreen-Rhizo automated platform for 24 days as previously described (48). For the MRI measurements, the seeds were placed in a Petri dish on wet filter paper. The Petri dish was sealed with parafilm and stored lightproof for 24 h in the growth chamber (16 °C/20 °C night/day temperature, 14 h light per day) to induce germination and only germinated seeds were used for further experiments. Seeds were subsequently sown in field soil (Sp2.1, Landwirtschaftliche Untersuchungs- und Forschungsanstalt, Speyer, Germany). Soil moisture was kept at 8.9%_m/m_, corresponding to 40% of the maximal water holding capacity (49). Per genotype, 18 seeds were planted in one pot (Ø=12.5 cm, 12 cm height) in a hexagonal grid with 2.5 cm seed spacing. Seedlings were imaged after 3 days in the growth chamber. For a longer experiment, single seeds were planted into larger pots (Ø=9 cm, 30 cm height) and were grown for one week before imaging.

Durum wheat (*Triticum turgidum*) *egt2* mutants were identified from a TILLING population developed in tetraploid cv Kronos (32). Two selected lines (Kronos2138 and Kronos3589) carrying premature termination codons in the two *EGT2* homoeologous coding sequences (TraesCS5A01G102000 and TraesCS5B02G164200LC) were crossed and F_1_ plants were self-pollinated. Progenies of selected wild-type and double mutant F_2_ individuals derived from two independent initial crosses were grown in rhizoboxes for seminal root angle analysis. Seeds were washed in 70% ethanol for 1 min, then in 1% sodium hypochlorite + 0.02% TritonX-100 for 5 min and rinsed with distilled water. Sterilized seeds were pre-germinated for 24 h at 28 °C in wet filter paper. Only germinating seeds were transferred in rhizoboxes for 7 days at 25 °C.

### Phenotyping experiments and rotation tests

For analysis of the root angles, plants were grown in rhizoboxes for 7 (seminal root angle) or 14 days (lateral root angle). The seminal root angle was measured as angle from the shoot to the root tip, in relation to the horizontal. For the angle of the lateral roots, the angle was measured from the outgrowth point of the main root to the lateral root tip in comparison to the horizontal. 20 randomly chosen lateral roots were measured per plant. For the rotation tests, the plants were grown in rhizoboxes for 5 days and then rotated once by 90°. For phytohormone treatments, the plants were grown for 5 days on agar without phytohormones and then transferred to agar plates supplemented with phytohormones as indicated in the results. After 1 h recovery, the agar plates were rotated once by 90°. Pictures were taken at the time points indicated in the graphs. For analysis, the root angle of every single root tip was measured in relation to the horizontal and the angle right after rotation was set to 0. For all measurements, the average of all roots per plant was calculated, presented in the graphs and compared in the statistical tests. For analysis of growth in the rhizotrons, root images were collected every two days, enabling to distinguish between seminal and crown roots. Images at 24 days were utilized for seminal, nodal and lateral root angle analysis. Root angle values were collected with the software ImageJ (50).

### RNA *in situ* hybridizations

Probes for *EGT2* (HORVU5Hr1G027890) mRNA were prepared from the whole coding sequence (start to stop codon). Cloning and RNA probe synthesis was performed as described before (33). RNA *in situ* hybridizations on roots of 7-day-old plants was performed as described before (33).

### Protein localisation in *Nicotiana benthamiana*

Constructs for transient expression in tobacco (*N. benthamiana*) were built using the greengate system, with the *EGT2* (HORVU5Hr1G027890) CDS in the pGGC module, the 35S promoter in the pGGA module (pGGA004), the UBQ10 terminator (pGGE009) and hygromycin resistance in the pGGF005 vector (51). TagRFP or mVENUS were used as fluorophors (52), either in the pGGB module as N-tag or in the pGGD module as C-tag (Table S5). The *Agrobacterium tumefaciens* strain GV3101 was transformed with expression clones and cultured in lysogeny broth medium. Bacterial cultures were precipitated and dissolved in 10 mM MgCl2, 10 mM MES, and 100 μM AS and infiltrated into tobacco leaves. Transgene expression was analyzed 3 days after infiltration.

### Bulked segregant analysis (BSA) and whole genome sequencing (WGS)

BSA (53) was carried out using plants from an F_2_-population obtained starting from the cross TM2835 × cv. Barke and segregating for the *EGT2* locus. 106 F_2_-seedlings were grown in flat rhizoboxes composed by two black plastic panels of 38.5 × 42.5 cm. Five pre-germinated seeds (1 day, 20 °C, on wet filter paper, in the dark) were positioned between moist filter paper sheets within each rhizobox. Each rhizobox was placed vertically in a larger plastic tank containing deionized water to a level of 3 cm from the bottom, in growth chamber) at 18 °C at night (8 h) and 22 °C at day (16 h) for 13 days. At the end of the growing period, root growth angle of seedlings was visually evaluated and a segregation rate of 88:18 (wild-type vs. hypergravitropic) recorded confirming that *egt2* segregates as a monogenic recessive Mendelian locus (χ^2^ 3:1 = ns) as previously described (25). Immediately after this inspection, 15 plants showing wild-type root growth angle and 15 plants showing an hypergravitropic angle were chosen for DNA preparation on a single plant basis using 2 cm^2^ leaf portions as previously described (45). DNA samples for BSA were obtained by mixing equal DNA amounts of each of the 15 bulk components, to a final concentration of 50 ng/ul. The two DNA bulks (in double) along with single plant DNA samples from 10 hypergravitropic plants were genotyped using the 9k Illumina Infinium iSelect barley SNP array (54). SNPs signal was analyzed using GenomeStudio (Illumina, San Diego, Inc.). For DNA bulks, SNPs signal was interpreted using the theta value approach as described in (55), modified in order to integrate for each SNP the signals obtained from two bulks (wild type or +/+ and hypergravitropic or −/−) in the “delta theta” value score as follows “delta theta” = [(theta bulk+/+)-(theta bulk −/−)]^2^.

Genomic DNA of TM2835 for whole genome shot gun sequencing was extracted from leaf samples using a commercial kit (Macheray-Nagel Nucleospin® Plant II). The DNA was sequenced with Illumina HiSeq PE150, and 699,353,963 paired-end reads were produced corresponding to a coverage of approx. 40×. Reads were aligned to the first version of barley cv. Morex reference genome (24) with BWA v.7.12 (56) and variants in the genomic space were called with SAMtools v. 1.3 (57, 58), filtering for a minimum reads depth of 5×, PHRED quality > 40. In order to discard background mutations due to the differences between the official Morex reference and the Morex parental seeds which had previously been used in the mutagenesis, the SNP calling considered further eight TILLMore mutants WGS data that was available at that moment, filtering with a custom AWK script for a minimum ratio DV/DP of 0.8 for the *egt2* mutant and a maximum ratio of 0.2 in every other mutant, where DP is the coverage depth at the SNP position and DV is number of non-reference bases at the same position. SNP effects were predicted with SNPEff v.3.0.7 (59).

For coverage analysis, a minimum of 5× read depth was considered, resulting in a target region of 3.5 GB containing a total of 15,805 mutations, hence the estimated mutation load on the entire genome is 22,579 mutations, or approx. 1 mutation per 220 kb which is of the same order of magnitude of mutation density (1 per 374 kb) formerly estimated based on TILLING results from the same TILLMore population (24). For the provean analysis, values <−2.5 were considered deleterious and values >−2.5 were tolerated.

### Modified pseudo-Schiff propidium iodide (mPS-PI) and Lugol staining

The mPS-PI staining was performed as described in (33). For Lugol staining, roots were fixed in 4% para-formaldehyde in phosphate buffered-saline (PBS) over night, embedded in 13% agarose and sectioned at the vibratome with 80 μm thickness. Then they were stained with Lugol solution for 5 min and rinsed with water.

### Microscopy

Transient expression in tobacco leaves was examined with a 25x Zeiss water-immersion objective using a Zeiss LSM 780 confocal microscopy system. tagRFP was excited with a 543 nm using a Helium Neon laser with emission detection through the meta-channel at 579 to 633 nm, the laser power is 30%. mVenus was excited at 488 nm by the argon laser with a 2.6% laser power, and emission was detected at 517 to 553 nm via the meta- channel (ChS2). DAPI was excited with a 405 nm using a diode laser, the laser power is 16%, and emission was detected at 440-480 nm. RNA *in situ* hybridization and Lugol-stained samples were examined using a Zeiss PALM MicroBeam microscope.

### MRI

MRI images were acquired on a 4.7 T vertical magnet equipped with a Varian console (60). A multi slice spin echo sequence was used. Sequence parameters were adapted to the different pot sizes. For the 9 cm pots, we used a birdcage RF coil with 10 cm diameter and the following sequence parameters: 0.5 mm resolution, 1 mm slice thickness, 9.6 cm field of view, TE = 9 ms, TR = 2.85 s, Bandwidth = 156 kHz, 2 averages. For the 12.5 cm pots, following parameters were changed: birdcage RF coil with 140 mm diameter, 14 cm field of view, and 0.55 mm resolution.

### CRISPR

For CRISPR target sequences, we choose twenty base pair sequences with the protospacer adjacent motif PAM sequence NGG in the first exon of *EGT2* (HORVU5Hr1G027890) that we checked at http://crispr.dbcls.jp/ for off-targets in the barley genome (Barley (*Hordeum vulgare*) genome, 082214v1 (Mar, 2012)). We used sites with only one predicted target for a 20mer and only up to 3 predicted targets for the 12mer target sequence upstream of the PAM. The CRISPR guide sequences are marked in Supplementary Figure 4A. The sgRNA shuttle vectors pMGE625 and 627 were used to generate the binary vector pMGE599 as described in (61). Transformation was carried out with the spring barley cv. Golden Promise grown in a climate chamber at 18 °C / 14 °C (light/dark) with 65% relative humidity, with a 16 h photoperiod and a photon flux density of 240 μmol m-^2^ sec-^1^. The binary vector pMGE599 was introduced into *Agrobacterium tumefaciens* AGL-1 strain (62) through electroporation (*E. coli* Pulser; Bio-Rad, http://www.bio-rad.com). The scutella tissue of barley caryopsis was used for *Agrobacterium*-mediated transformation as described previously (63). The insert integration in the barley genome was confirmed by detection of hygromycin gene sequences via PCR in generated T0 lines and were analyzed for mutations in *EGT2* by PCR and Sanger sequencing and the seeds for T1 generation were used for experiments (Table S5).

### qRT-PCR

For the qRT-PCR, RNA from plants grown for 7 days after germination in rhizoboxes was extracted with the RNeasy Plant Mini Kit (Qiagen) and first strand cDNA was synthesised with the RevertAid First Strand cDNA synthesis Kit (ThermoFisher). Ca. 12 plants were pooled for one biological replicate per sample and samples were divided into root tip (containing meristem and elongation zone, ~2 mm until the outgrowth of root hairs) and adjacent differentiation zone (~8 mm). For each genotype, 4 biological replicates and 3 technical replicates were used. For the reaction, 2 μl of PerfeCTa SYBR Green SuperMix (Quantabio), 1 μl primer mix of a concentration of 1 μM and 1 μl cDNA was mixed. The primer efficiency of each oligonucleotide was calculated using the following dilution series: 1, 1/2, 1/4, 1/8, 1/16, 1/32, 1/64, and 1/128. The relative expression levels of the transcripts were calculated with reference to the housekeeping gene tubulin (HORVU1Hr1G081280) and according to the method described in (64). Significant differences in gene expression levels were determined by a two-sided Student’s t-test.

### Laser capture dissection microscopy (LCM) and RNAseq

Root tips of the most vertically grown seminal root of 3-day-old plants were used and assigned as one biological replicate. Per genotype, 4 biological replicates were analyzed. Plants were grown in rhizoboxes and fixed with Farmers fixative (EtOH:Acetic acid 3:1) on ice for 15 min under 500 mbar vacuum and subsequent swirling at 4 °C for 1 h. The fixation solution was replaced and the procedure was repeated twice before replacing the solution with 34% sucrose and 0.01% safranin-O in PBS. The samples were vacuum-infiltrated again for 45 min and incubated on ice at 4 °C for 21 h. Then the samples were dried carefully with tissue paper and embedded in tissue-freezing medium as described before (65). The medium blocks containing the tissue were stored at −80 °C and were acclimatized to −20 °C in the cryomicrotome (Leica CM1850). Longitudinal sections of 20 μm thickness were mounted on poly-L-lysine-coated glass slides (Zeiss) and the tissue-freezing medium was removed after 5:30 min by incubation in 50% EtOH and 1 min incubation each in 70% EtOH, 95% EtOH, 100% EtOH and 100% xylol (RNase-free). The tissues (root cap, meristem and 900 μm of the elongation zone adjacent to the meristem) were cut with the following settings of the PALM Microbeam laser capture instrument (Zeiss, Germany): Energy: 79, speed: 100, Cutting Program: “Center RoboLPC”, picked up manually with a sharp needle and transferred to the cap of RNAse-free adhesive caps (Zeiss). RNA was isolated with the Arcturus PicoPure RNA Isolation Kit (Thermo Fisher) according to the manufacturer’s protocol for tissue, including the DNase treatment. RNA quality was determined with an Agilent 2100 Bioanalyzer using the Agilent RNA 6000 Pico kit and yielded RIN values between 7.1 and 8.9 and a concentration between 610 and 95.000 pg/μl (Table S3). Pre-amplification and library preparation was carried out as described in (66). Detection and sequenced on an Illumina NovaSeq sequencing instrument with a PE100 protocol. The RNA-Seq experiments yielded on average 41 million 100 bp paired-end reads per sample (Table S3). Trimmomatic version 0.39 (67) was used to remove low-quality reads and remaining adapter sequences from each read dataset. Specifically, a sliding window approach was used, in which a read was clipped if the average quality in a window of 4 bp fell below a phred quality score of 15. Only reads with a length of ≥30☐bp were retained for further analyses. Data are deposited at the sequence read archive (SRA), PRJNA589222. BBDuk of the BBTools suite (https://jgi.doe.gov/data-and-tools/bbtools/) was employed to remove rRNA reads from the datasets using a kmer length of 27 as filtering threshold for decontamination. After removal of rRNA reads, on average 8 million paired reads remained. The splice-aware STAR aligner v.2.7.2b (68) was used to align the remaining reads against a genome index of the barley reference sequence and annotation of genotype Morex (IBSC v2.0) (24). Multi-mapping reads that mapped to more than one position were excluded from subsequent steps by considering only reads, which mapped in a single location (--outFilterMultimapNmax 1). On average 5 million reads per sample aligned to unique positions in the gene set of the IBSC v2.0 barley reference genome with 46,272 predicted coding and non-coding gene models (EnsemblPlants release 45, (24), Table S3). The aligned paired-end reads were ordered according to their position and transformed to .bam files by the software samtools (version 1.3.1, (57)). Alignment of sequences to the reference genome of Morex (release 45) (24) was performed using HTSeq (version 0.10.0, (69)) with the parameters ‘-r pos -i gene_id -s no --secondary-alignments ignore --supplementary-alignments ignore’. The principal component analysis (PCA) was performed on the expression data using the normalization procedure rlog() implemented in the R package DESeq2 and the plotPCA() function (version 1.22.2, (34)). Expression values were normalized with library size by calculating fragments per million (FPM) reads using the fpm() function of DESeq2, after removal of lowly expressed genes with less than 10 reads over all samples. Expression levels of genes were estimated by the variance-mean dependence in the count table based on a generalized linear model using the negative binomial distribution within the R package DESeq2 (34) calculating log_2_ fold change (log_2_FC) values between wild type and mutant in the respective tissues with the design ~ genotype + tissue + genotype:tissue. Significance values for log_2_FC values were calculated as Wald test *p*-value and were adjusted by the Benjamini-Hochberg procedure to obtain false discovery rates (FDR) (70). Genes with a FDR <5% and log_2_FC of |1| were considered differentially expressed. From this gene set, we excluded gene pairs that were assigned to chr0 and chr1 which had the same annotation and the respective gene partner was the one with the closest related transcript after a BLAST search. Gene ontology term enrichment of the resulting gene set was performed using agriGo (71). The sequencing data have been deposited in the NCBI sequencing read archive (PRJNA589222).

### Phylogenetic analysis

The EGT2 (HORVU.MOREX.r2.5HG0370880.1) protein sequence was BLASTed on Phytozome v12.1 to the *Brachypodium dystachyon* proteome v3.1, the *Oryza sativa* proteome v7_JGI, the *Zea mays* proteome Ensembl-18, the *Arabidopsis thaliana* proteome TAIR10, the *Prunus persica* proteome v2.1 and the *Sorghum bicolor* proteome v3.1.1 with the default settings. Hits with E-values <3.9E-77 were considered. The identified orthologs were then confirmed using the EnsemblPlants Compara Ortholog tool. Retrieved protein sequences were aligned by ClustalW in the software MEGA X, with default values (72): Ancestral states were inferred using the Maximum Likelihood method (73) and JTT matrix-based model (74). The tree shows a set of possible amino acids (states) at each ancestral node based on their inferred likelihood at site 1. The initial tree was inferred automatically by applying Neighbor-Join and BioNJ algorithms to a matrix of pairwise distances estimated using the JTT model, and then selecting the topology with superior log likelihood value. The rates among sites were treated as being uniform among sites (Uniform rates option). This analysis involved 16 amino acid sequences.

## Acknowledgments

This work was funded by the Deutsche Forschungsgemeinschaft (DFG) grant HO2249/21-1 to FH. KAN, DF and RK acknowledge support from the Helmholtz Association for the Forschungszentrum Jülich. The rhizotron study received funding from the European Union’s Horizon 2020 research and innovation programme under grant agreement No 731013 (EPPN2020). Work described here is supported in part by the project ‘Rooty- A root ideotype toolbox to support improved wheat yields’ funded by the IWYP Consortium (project IWYP122) to CU, JS, RT and SS, via the Biotechnology and Biological Sciences Research Council in the United Kingdom (BB/S012826/1).

The authors thank Dr. Felix Frey (University of Bonn) for his advice on RNAseq data analysis and discussion, Dr. Johannes Stuttmann (University of Halle) for sharing the CRISPR/Cas cloning vectors and Shalima H. Orse for support throughout the project. We thank Anna Galinski, Jonas Lentz, Carmen Müller, Bernd Kastenholz, Ann-Katrin Kleinert, Roberta Rossi, and Kwabena Agyei (Forschungszentrum Jülich GmbH) for their assistance during the rhizotron study. RK and DP gratefully acknowledge Dagmar van Dusschoten and Johannes Kochs for support and maintenance of the MRI System.

---

**Supplementary Figure 1: Root phenotype of *egt2-1***

**A** Wild type and *egt2-1* roots grown on paper sheets, 14 days after germination (DAG). Scale bar: 2 cm.

**B** Number of seminal roots 7 DAG; n= 40 per genotype in one experiment.

**C** Root length 14 DAG; n = 8-9 per genotype in two independent experiments.

**D** Root length after rotation; n = 32 in three independent experiments; compare Figure 1G; standard deviation is depicted; two-tailed t-test did not show any significant differences between the genotypes at respective time points; all measurements were normalized to the starting length of the roots at time point 0.

**E** Magnetic resonance imaging (MRI) pictures of wild type and *egt2-1* roots grown in soil for 7 DAG. Scale bar: 2 cm.

**F** Root angle of roots grown in soil for 3 DAG and captured by MRI (see D); n = 17-18 per genotype; two-tailed t-test, ** *p* <0.01.

**G** Root angle of seminal roots of plants grown in rhizotrons, measured by angle between the outermost seminal roots; 26 DAG; n = 10-11 per genotype; two-tailed t-test, ** *p* <0.01.

**H** Root angle of lateral roots of plants grown in rhizotrons, angle measured between main roots and lateral roots; 26 DAG: n = 5 (30 lateral roots per plant) per genotype; two-tailed t-test, ** *p* <0.01.

**Supplementary Figure 2: Meristem phenotype of *egt2-1* resembles the wild type phenotype**

**A** Root meristem of wild type and *egt2-1* 7 DAG; arrow heads mark transition to elongation zone. Scale bar: 200 μm.

**B** Wild type and *egt2-1* root cap 7 DAG. Scale bar: 50 μm; brightness adjusted in magnification.

**C** Quantification of meristem length 7 DAG; n= 15-20 roots per genotype; two-tailed t-test does not show any significant difference.

**D** Lateral roots of wild type and *egt2-1* plants 14 DAG stained by mPS-PI staining. Scale bar: 50 μm.

**Supplementary Figure 3: The *egt2-1* mutant and the corresponding wild type react similarly to auxin treatment**

**A**, **C**, **E**, **G**, **I** Angle of the root tip after indicated treatment and rotation for 90° (time point 0).

**B**, **D**, **F**, **H**, **J** Length of the roots after indicated treatment and rotation for 90° (time point 0); the two genotypes were compared between each other at the respective time points by a two-tailed t-test, * *p* < 0.05; standard deviation is depicted; to account for the different starting angles of the roots, all measurements were normalized to the starting angle/length of the roots at time point 0.

**A**, **B** IAA treatment; n = 4-6 per genotype and treatment in two independent experiments.

**C**, **D** NAA treatment; n = 4-6 per genotype and treatment in two independent experiments.

**E**, **F** 2,4D treatment; n = 5-10 per genotype and treatment in three independent experiments.

**G**, **H** NPA treatment; n = 7-10 per genotype and treatment in three independent experiments.

**I**, **J** Cytokinin treatment; n = 4-8 per genotype and treatment in two independent experiments

**Supplementary Figure 4: CRISPR/Cas9 induced mutation in *EGT2* and conserved function in wheat**

**A** Gene model of *EGT2* and partial DNA sequence; arrow marks translation start site; purple boxes mark the CRISPR target sites; in the *egt2-2* mutant line, CRISPR/Cas induced a deletion between the target sites as depicted.

**B** Protein structure of EGT2 with the SAM domain from amino acid 172 - 225; protein alignment of EGT2, AtSAM5 (At3g07760) and peach WEEP (Prupe. 3G200700) in the SAM region.

**C** Root length 7 DAG, two-tailed t-test does not show a significant difference (*p* <0.05); n = 15-17 in two independent experiments.

**D** Wild type GP and *egt2-2*lateral roots 14 DAG. Scale bar: 1 cm.

**E** Root tip angle after rotation; plants 5 DAG were rotated by 90° (time 0) and the root tip angle was measured over time; n = 20 per genotype in two independent experiments; the two genotypes were compared between each other at the respective time points by a two-tailed t-test and no significant difference was detected; standard deviation is depicted; to account for the different starting angles of the roots, all measurements were normalized to the starting angle of the roots at time 0.

**F** Root length after rotation as described in D; n = 20 per genotype in two independent experiments.

**G** Area enclosed by the first three seminal root of wild type (WT/WT) and *egt2* (mut/mut) wheat seedling at 7 DAG; n = 16 and 35 for wild type and mutant, respectively.

**H** Number of seminal roots of wild type (WT/WT) and *egt2* (mut/mut) wheat seedling at 7 DAG; n = 18 and 39 for wild type and mutant, respectively.

**I** Length of the first three seminal root in wild type (WT/WT) and *egt2* (mut/mut) wheat seedling at 7DAG; n = 16 and 35 for wt and mutant, respectively.

Wheat plants analyzed in G, H and I were derived from two independent segregating populations; the two genotypes were compared by a two-tailed t-test (* p <0.05, ns = no significant difference).

**Supplementary Figure 5: Phylogenetic tree of EGT2 and related proteins**

Species abbreviations: AT: *Arabidopsis thaliana*; BGIOS: *Oryza sativa Indica Group*; BRADI: *Brachypodium dystachyon*; Os: *Oryza sativa Japonica Group;* PRUPE: *Prunus persica*; SORBI: *Sorghum bicolor;* Traes: *Triticum aestivum;* TRITD: *Triticum durum;* Zm: *Zea mays.*

**Supplementary Figure 6: Heat map of differentially expressed genes (FDR <5% and log2FC >|2|) between wild-type and egt2-1 in root cap, root meristem and elongation zone**

Average of log1p of fpm normalized counts of four biological replicates of the respective genotype and tissue is shown in blue scale.

**Supplementary Figure 7: Expression of expansins**

Fpm normalized expression values of the expansins differentially expressed in the gene set; padj is depicted with * *p* <0.05, ** *p* <0.01; RC: root cap, M: meristem, EZ: elongation zone.

**Supplementary Table 1. BSA mapping results**

**Supplementary Table 2. SNPs whole genome sequencing**

**Supplementary Table 3. Overview of RNAseq reads and mapping results**

**Supplementary Table 4: DEGs**

**Supplementary Table 5: primer sequences**

