## Supplementary figures and images for "*ENHANCED GRAVITROPISM 2* encodes a STERILE ALPHA MOTIVE containing protein that controls root growth angle in barley and wheat"

### Supplementary Figure 1

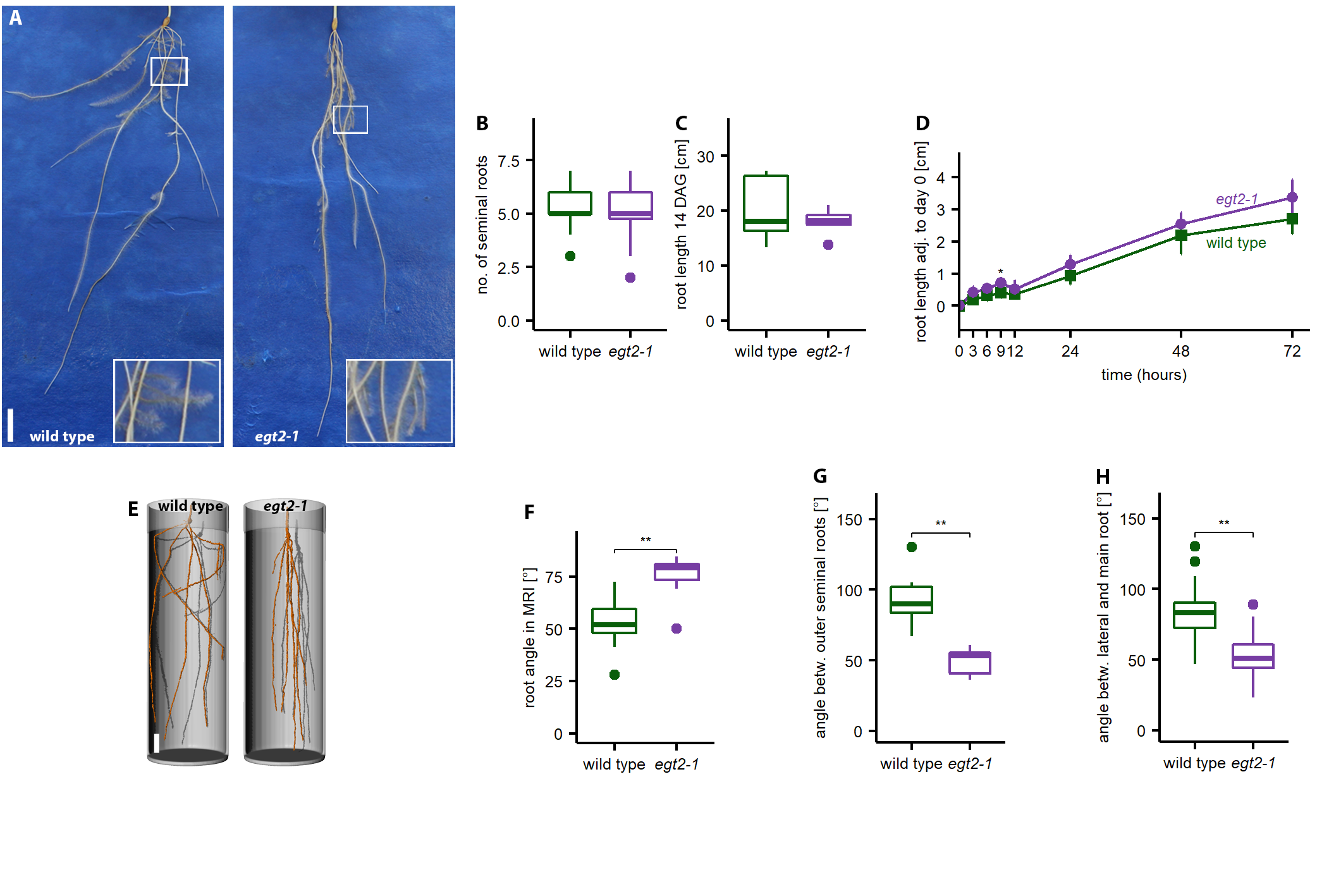

### Supplementary Figure 2

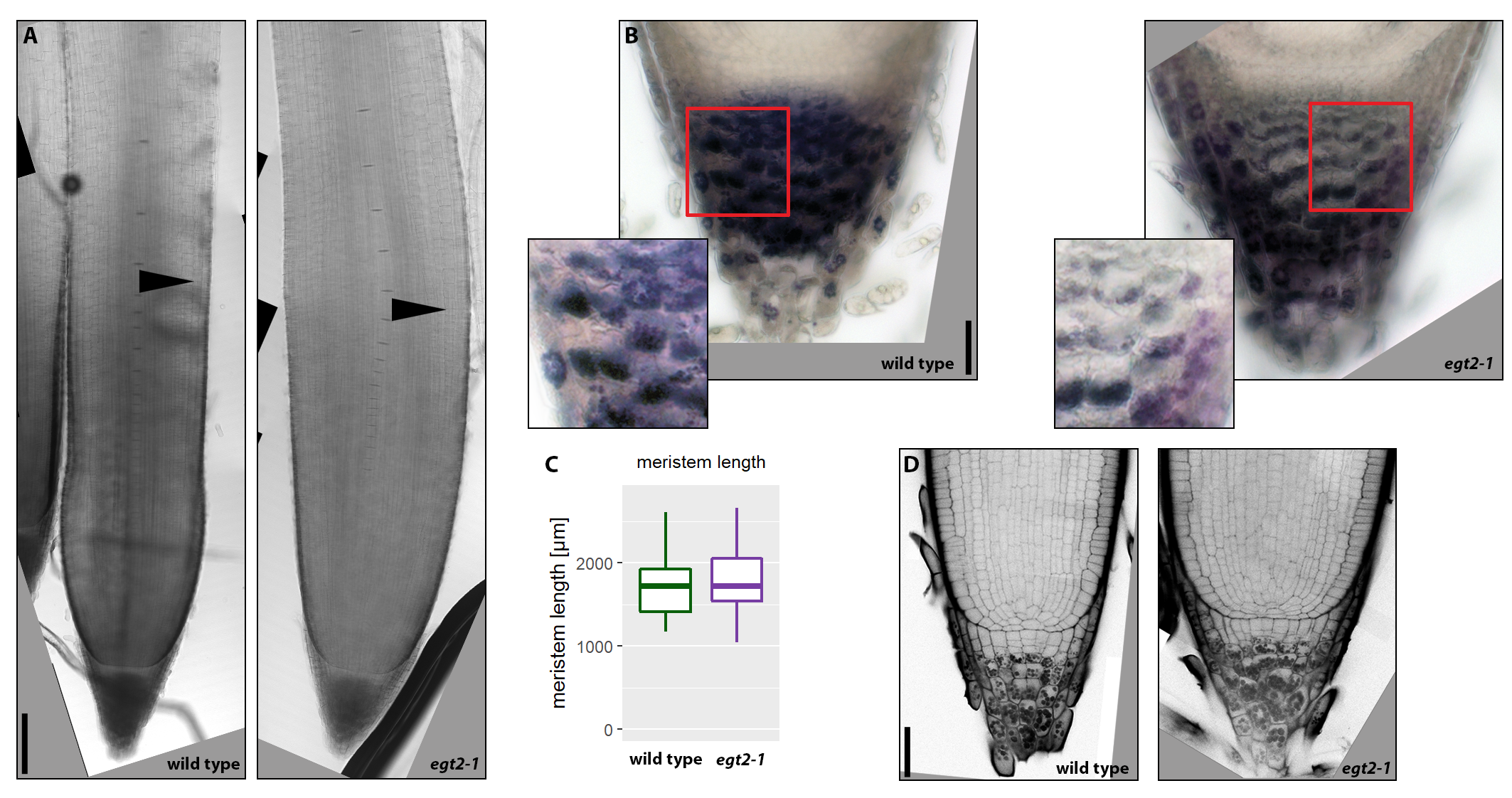

### Supplementary Figure 3

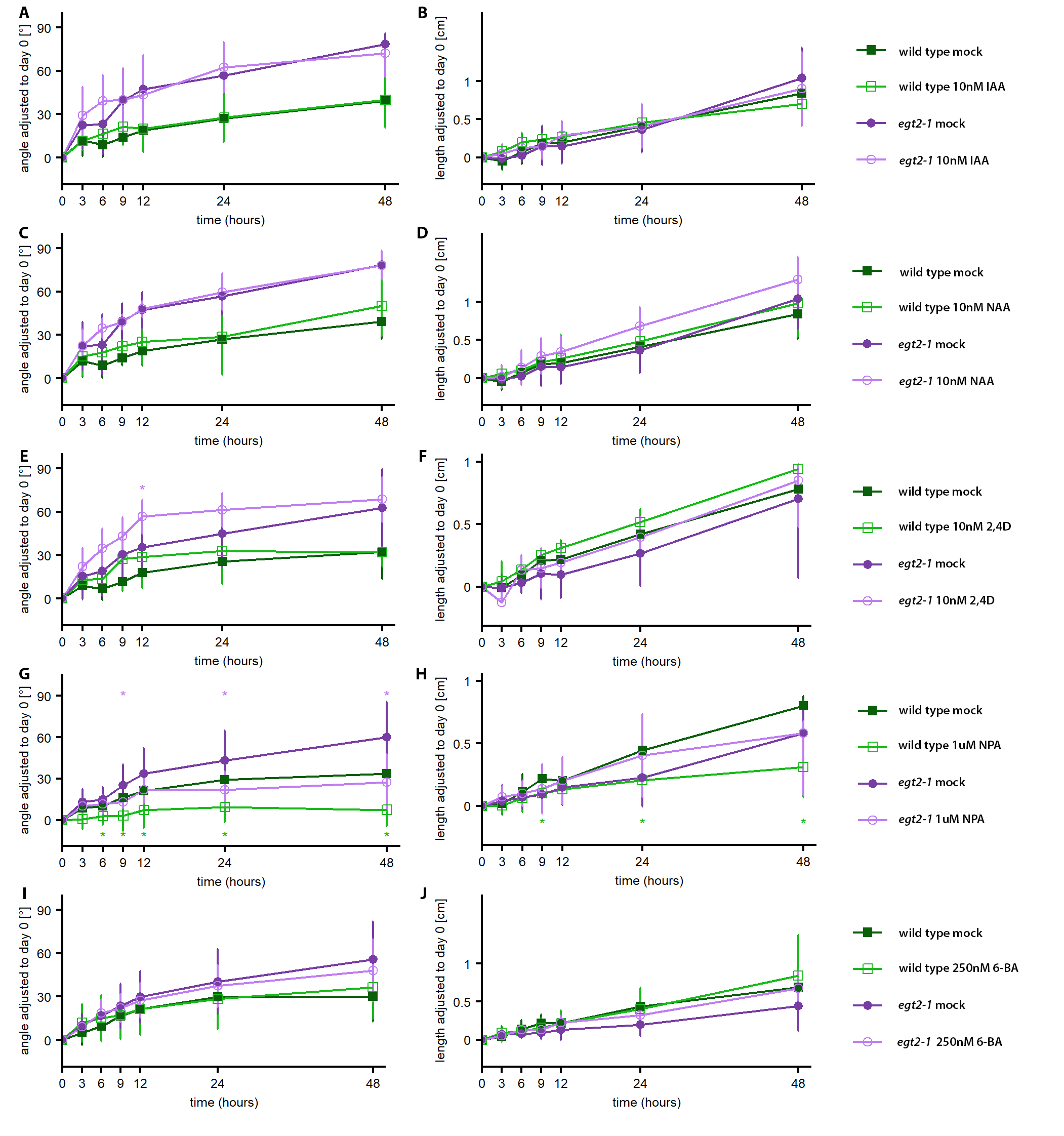

### Supplementary Figure 4

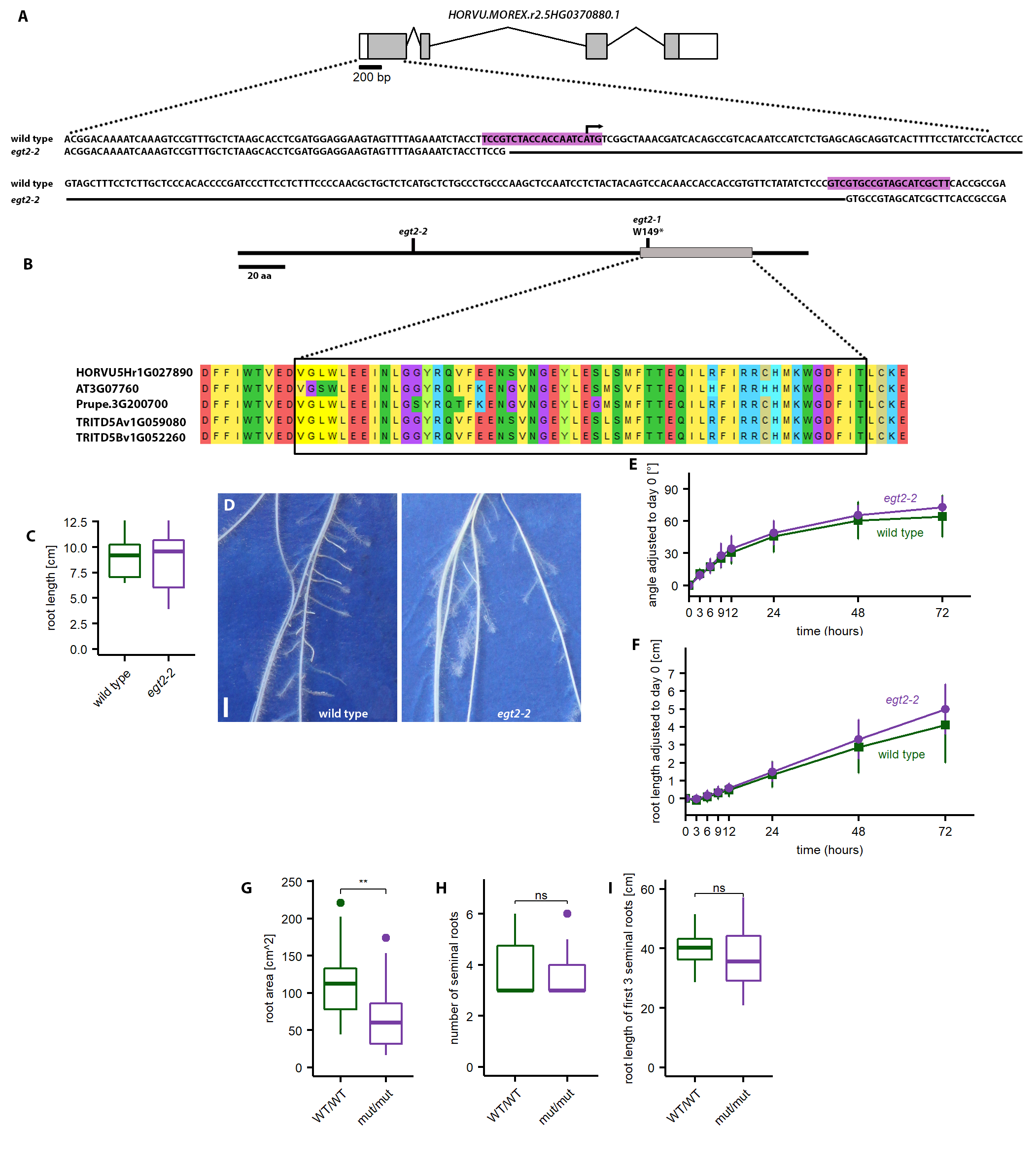

### Supplementary Figure 5

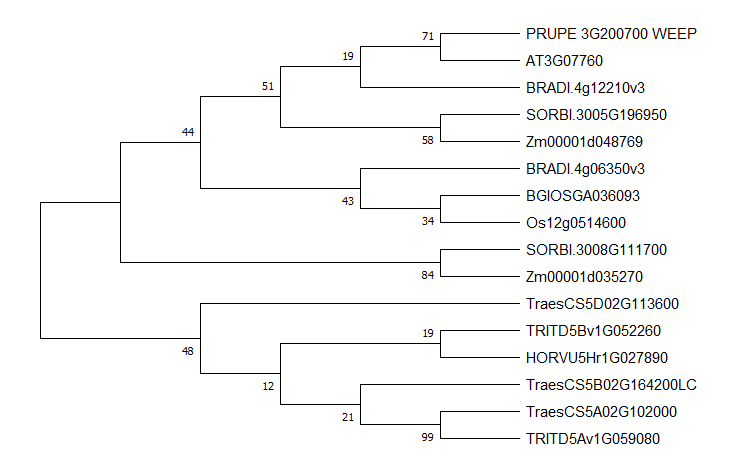

### Supplementary Figure 6a

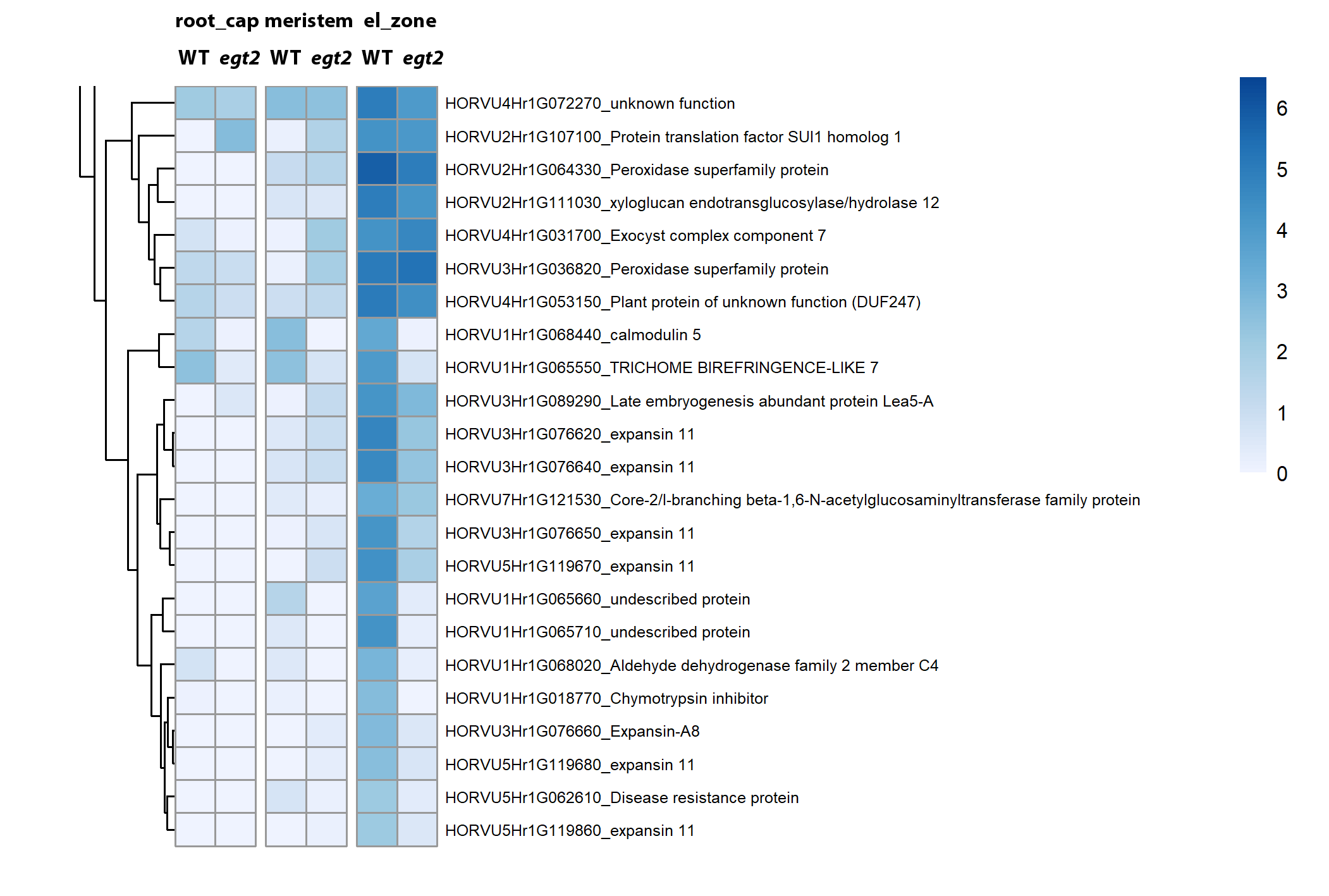

### Supplementary Figure 6b

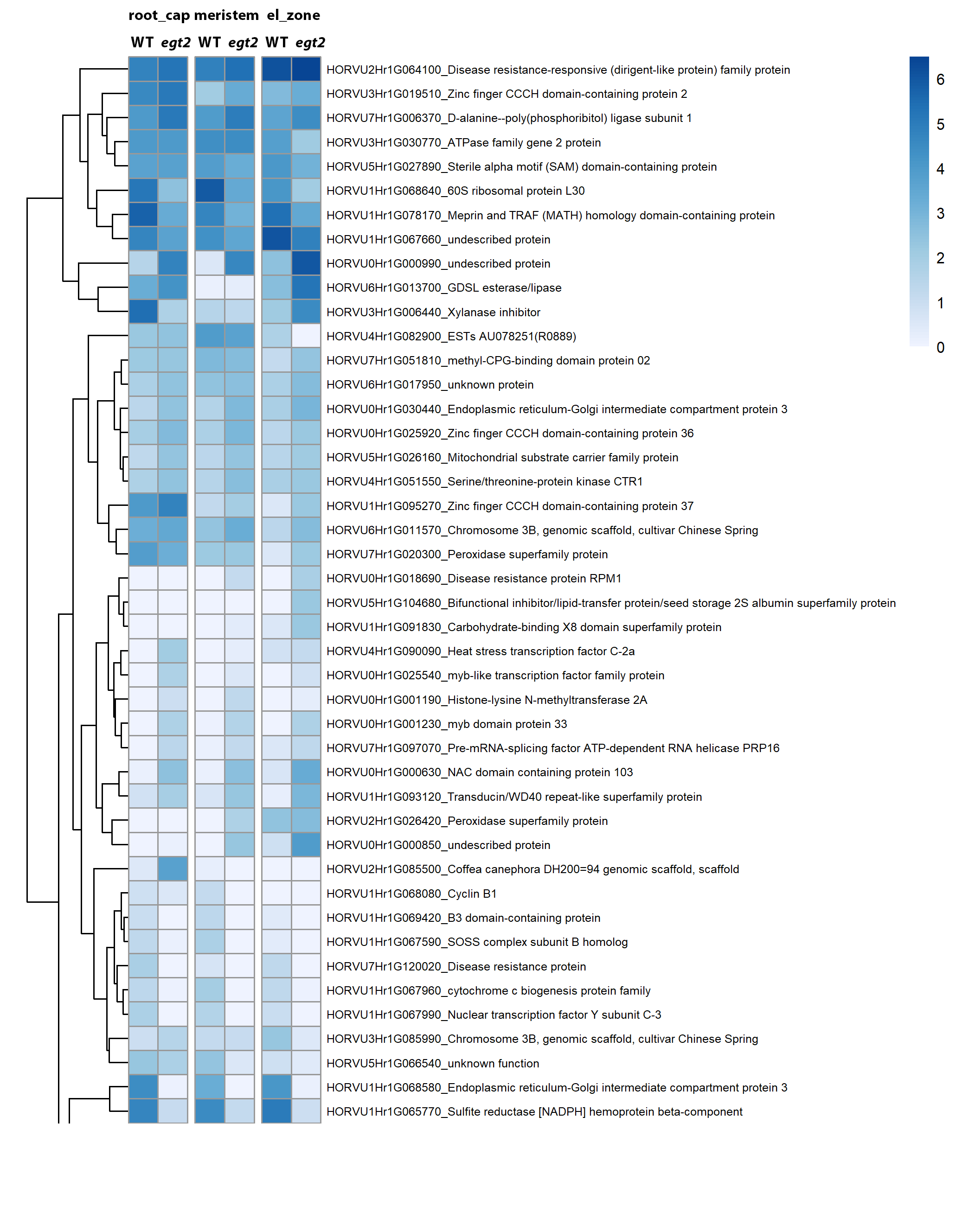

### Supplementary Figure 7

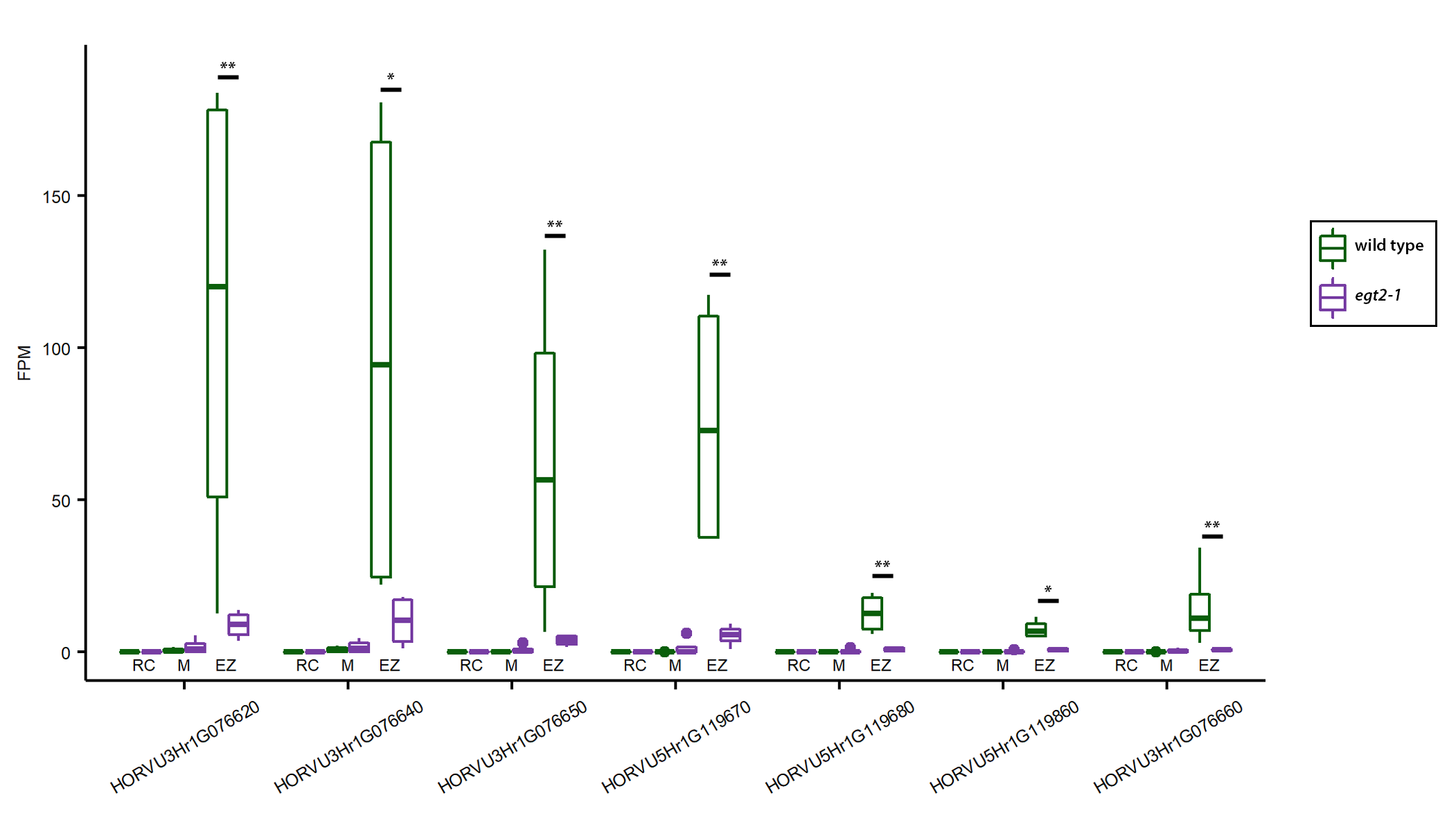
